# OligoPVP: Phenotype-driven analysis of individual genomic information to prioritize oligogenic disease variants

**DOI:** 10.1101/311654

**Authors:** Imane Boudellioua, Maxat Kulmanov, Paul N Schofield, Georgios V Gkoutos, Robert Hoehndorf

## Abstract

**Purpose:** An increasing number of Mendelian disorders have been identified for which two or more variants in one or more genes are required to cause the disease, or significantly modify its severity or phenotype. It is difficult to discover such interactions using existing approaches. The purpose of our work is to develop and evaluate a system that can identify combinations of variants underlying oligogenic diseases in individual whole exome or whole genome sequences.

**Methods:** Information that links patient phenotypes to databases of gene–phenotype associations observed in clinical research can provide useful information and improve variant prioritization for Mendelian diseases. Additionally, background knowledge about interactions between genes can be utilized to guide and restrict the selection of candidate disease modules.

**Results:** We developed OligoPVP, an algorithm that can be used to identify variants in oligogenic diseases and their interactions, using whole exome or whole genome sequences together with patient phenotypes as input. We demonstrate that OligoPVP has significantly improved performance when compared to state of the art pathogenicity detection methods.

**Conclusions:** Our results show that OligoPVP can efficiently detect oligogenic interactions using a phenotype-driven approach and identify etiologically important variants in whole genomes.

## Introduction

Discrimination of causative genetic variants responsible for disease is a major challenge. An increasingly large family of algorithms and strategies has been developed to aid in identification of such variants^1^. These methods use properties of variants such as evolutionary conservation, predicted structural changes, allele frequency and function to predict pathogenicity. For variants in non-coding sequence regions, additional information used by computational models includes predicted regulatory function and recognized DNA–protein or DNA–RNA interactions^1–3^. Furthermore, phenotype annotations to human and model organism genes can be added to provide another layer of discrimination between involved pathogenic and non-pathogenic variants^4–6^. Phenotype-based methods can identify the likelihood that a particular gene or gene product may give rise to phenotypes observed in an individual^7,8^.

The increasing availability of patient sequence information coupled with resources that provide a detailed phenotypic characterization of diseases, as well as the wealth of gene-to-phenotype associations from non-human disease models^9^, are now enabling new approaches to the prioritization of causative variants and facilitating our ability to dissect the genetic underpinnings of disease^5^. PhenomeNET^10^, developed in 2011, is a computational framework that utilizes pan-phenomic data from human and non-human model organisms to prioritize candidate genes in genetically-based diseases^10^. We have combined PhenomeNET with genome-wide pathogenicity predictions to develop the PhenomeNET Variant Predictor (PVP)^4^ as a system that combines information about pathogenicity of variants with known gene–phenotype associations to predict causative variants. We recently developed the PVP system to classify variants into causative and non-causative^4^.

While PVP has a significantly better performance in the prioritization of single variants in monogenic diseases than competing algorithms, many diseases that have been traditionally considered as monogenic are increasingly being understood within the context of complex inheritance and multifactorial disease phenotypes. Recent evidence for oligogenicity has been reported for amyotrophic lateral sclerosis^11^ where some pathogenic rare variants were observed to be present as heterozygotes, hypertrophic cardiomyopathy^12^, Parkinson’s disease^13^, cardiac septal defects^14^, and Hirschprung’s disease^15^. These phenomena have been known for many years^16^, but as the basis of more “monogenic” Mendelian diseases has been identified, the search for further interacting variants has proved difficult due to limited availability of genetic data and consequently insufficient statistical power. Furthermore, there is a lack of strategies applied to individual patients for detecting variants which might, for example, be hypomorphic, might be subject to haploid insufficiency (and therefore pathogenic when heterozygous), or which are common (i.e., have a high minor allele frequency within a population). Modifier gene variants may be either rare or common. The assumption that a modifier gene must be rare in a population depends on whether it is associated with a phenotype subject to negative selection, and is overrepresented in all or some phenotypic sub-populations of patients. Identification of common modifier variants similarly depends on whether they are overrepresented in the patient population, but this is much more difficult and may require large patient populations to determine^16^. The importance of oligogenicity is increasingly being recognized, with classic cases being the recognition of Bardet-Biedl syndrome^17^ and Huntington’s disease^18^ as primarily oligogenic.

In most of the examples above, discovery of oligogenicity involved targeted examination of genes already known to be involved in the disease under investigation, for example through disease panels. Often, the selection of genes to include in such a panel relies on the availability of additional information about molecular or functional connections between the entities (genes or gene products) bearing the variants. Computational discovery of causative variants that are involved in oligogenic and polygenic diseases, in particular in genes not previously associated with the disease, is particularly challenging; such methods would have to be able to incorporate and utilize a large amount of background information about molecular and (patho-)physiological interactions within an organism. The observation that disease-implicated proteins often interact with each other has stimulated the development of network-based approaches to identification of disease modules. However, relevant interactions may occur across much larger distances within pathways and networks, or at the whole organism physiological level where systems knowledge is critical for understanding^19,20^. Phenotypes provide a readout for all of these disease-relevant interactions and offer insights into the underlying pathobiological mechanisms^21^.

Phenotype data can be a powerful source of information for variant prioritisation and is complementary to pathogenicity prediction methods based on molecular information. In a recent study, we were able to identify multiple variants in known thyroid disease genes in individual patients with congenital hypothyroidism^4^ as well as compound heterozygosity. It is an open question whether it is possible to use phenotype similarity to facilitate the computational discovery of multiple contributing variants in oligogenic diseases, or for identifying modifying variants that affect the pattern, severity, or onset of a disease.

Here, we first evaluate the success of PVP in identifying oligogenic combinations of variants. We then present OligoPVP, a novel algorithm for finding oligogenic combinations of variants in personal genomes. We apply OligoPVP to the simplest form of oligogenic inheritance, digenic inheritance, where mutations in two separate genes present in a single individual lead to a particular phenotypic manifestation that is not apparent in individuals carrying only one of these variations^22,23^. We show that OligoPVP can be used to identify gene variants in oligogenic diseases, and their interactions, and evaluate these on a set of synthetic whole genome sequences into which we insert multiple variants that together are causative for a complex disease. OligoPVP is freely available at https://github.com/bio–ontology–research–group/phenomenet–vp.

## Materials and Methods

### Digenic disease

The Digenic Disease Database (DIDA) v2^22^ consists of 258 curated digenic combinations representing 54 diseases, with 448 variants in 169 genes. Of the 258 digenic combinations, 189 have HPO annotations, representing 52 diseases, 153 distinct genes, and 337 unique variants. We use the 189 digenic combinations with HPO annotations in our experiments. 25 of these combinations are triallelic and exhibit compound heterozygosity in one gene while the remaining 164 combinations are biallelic.

We use the combinations of variants from DIDA to generate 189 synthetic whole genome sequences by randomly inserting the causative variants in a randomly selected whole genome sequence from the 1000 Genomes Project^24^.

### Interaction data

We downloaded all interactions occurring in humans from the STRING database version 105^25^. Then, we mapped all interactions to their respective genes using the mapping file provided by STRING to generate 989,998 interactions between genes, representing 13,770 unique genes. We use these interactions between genes to prioritize combinations of variants in OligoPVP.

### PhenomeNET Variant Predictor

The PVP system used in our analysis, the synthetic genome sequences we generated for the evaluation of our system, and our analysis results can be found at https://github.com/bio-ontology-research-group/phenomenet-vp.

## Results

### Prediction of di-allelic and tri-allelic disease variants

We analyze each WGS using the phenotypes provided for the combination of variants in DIDA. As complex diseases are often caused by combinations of variants that are common individually, we do not filter any variants by minor allele frequency. On average, each WGS in our experiments contains 2,192,967 variants.

We use the phenotypes associated with the combination of variants in DIDA as phenotypes associated with the synthetic WGS, and we use PVP^4^ to prioritize variants, using an “unknown” mode of inheritance model. Out of 164 whole genome sequences where two variants were inserted, we find both causative variants (i.e., the two variants we inserted) as the highest ranked variants in 88 cases (53.66%) and within the top ten ranks in 107 cases (65.24%) (see Table 1). For the 25 cases of triallelic diseases, we find all three causative variants within the first three ranks in 10 cases (40.00%) and we find all three causative variants within the top ten variants in 14 cases (56.00%) (see Table 2).

**Table 1.**
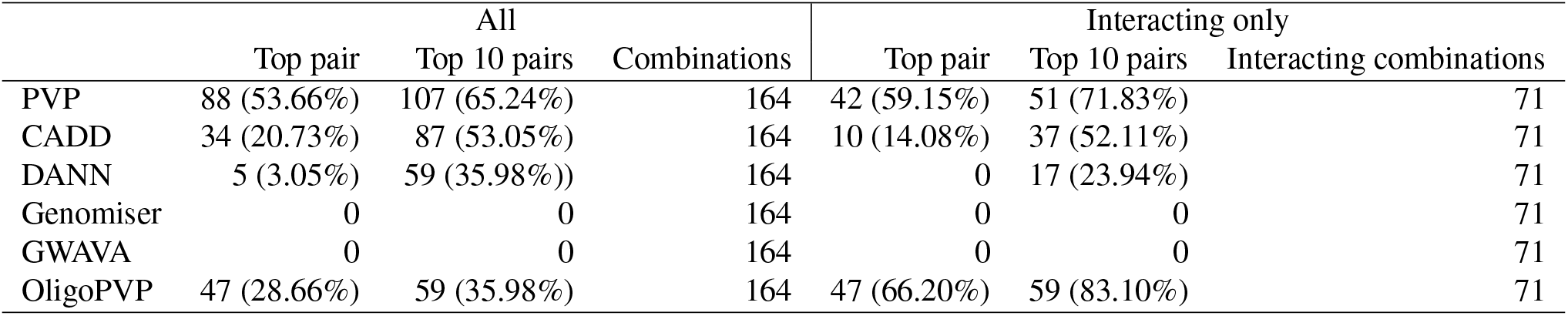
Comparison of different variant prioritization systems for recovering di-allelic variants. We split the evaluation in two parts, one in which we consider all variants and another in which we only consider variants for which we have background knowledge about their interactions.

|  | All |  |  | Interacting only |  |  |
| --- | --- | --- | --- | --- | --- | --- |
|  | Top pair | Top 10 pairs | Combinations | Top pair | Top 10 pairs | Interacting combinations |
| PVP | 88 (53.66%) | 107 (65.24%) | 164 | 42 (59.15%) | 51 (71.83%) | 71 |
| CADD | 34 (20.73%) | 87 (53.05%) | 164 | 10 (14.08%) | 37 (52.11%) | 71 |
| DANN | 5 (3.05%) | 59 (35.98%) | 164 | 0 | 17 (23.94%) | 71 |
| Genomiser | 0 | 0 | 164 | 0 | 0 | 71 |
| GWAVA | 0 | 0 | 164 | 0 | 0 | 71 |
| OligoPVP | 47 (28.66%) | 59 (35.98%) | 164 | 47 (66.20%) | 59 (83.10%) | 71 |

**Table 2.** Comparison of different variant prioritization systems for recovering tri-allelic variants. We split the evaluation in two parts, one in which we consider all variants and another in which we only consider variants for which we have background knowledge about their interactions.

|  | All |  |  | Interacting only |  |  |
| --- | --- | --- | --- | --- | --- | --- |
|  | Top triple | Top 10 triple | Combinations | Top triple | Top 10 triple | Interacting combinations |
| PVP | 10 (40.00%) | 14 (56.00%) | 25 | 7 (43.75%) | 10 (40.00%) | 16 |
| CADD | 4 (16.00%) | 9 (36.00%) | 25 | 7 (43.75%) | 12 (75.00%) | 16 |
| DANN | 0 | 6 (24.00%) | 25 | 0 | 4 (25.00%) | 16 |
| Genomiser | 0 | 0 | 25 | 0 | 0 | 16 |
| GWAVA | 0 | 0 | 25 | 0 | 0 | 16 |
| OligoPVP | 10 (40.00%) | 10 (40.00%) | 25 | 10 (62.50%) | 10 (62.50%) | 16 |

Individually, the performance of our approach differs significantly between diseases, depending on the availability of gene–phenotype associations and high quality and informative disease–phenotype associations in DIDA. Table 3 provides an overview of the performance of PVP for individual diseases, and we provide the full analysis results on our website.

**Table 3.** Analysis of top ranks of variants by PVP summarized by disease

|  | Top hit | Top 3 hits | Top 10 hits | Variants (Combinations) |
| --- | --- | --- | --- | --- |
| Familial long QT syndrome | 21 (50.00%) | 38 (90.48%) | 41 (97.62%) | 42 (21) |
| Kallmann syndrome | 18 (47.37%) | 27 (71.05%) | 27 (71.05%) | 38 (19) |
| Bardet-Biedl syndrome | 14 (36.84%) | 28 (73.68%) | 32 (84.21%) | 38 (14) |
| Alport syndrome | 14 (45.16%) | 28 (90.32%) | 29 (93.55%) | 31 (15) |
| Non-syndromic genetic deafness | 12 (50.00%) | 18 (75.00%) | 18 (75.00%) | 24 (12) |
| Oculocutaneous albinism | 6 (40.0%) | 12 (80.00%) | 12 (80.0%) | 15 (7) |
| Primary ovarian insufficiency | 2 (13.33%) | 2 (13.33%) | 2 (13.33%) | 15 (7) |
| Usher syndrome | 5 (33.33%) | 11 (73.33%) | 12 (80.0%) | 15 (7) |
| Hypodontia | 6 (50.0%) | 12 (100.0%) | 12 (100.0%) | 12 (6) |
| Others | 66 (38.15%) | 118 (68.21%) | 128 (73.99%) | 173 (81) |

In particular, for the case of hypodontia, PVP identifies all the causative variant pairs in the top 3 ranks in all synthetic patients, and in Familial long QT syndrome, the causitive variant pairs can be found in the top 3 ranks in over 90% of the synthetic patients. Similarly, for the case of Bardet-Biedl syndrome (BBS), PVP ranks 84.21% of causative variant pairs in the top 10, and identifies digenic causative variants in 9 of the 16-20 genes now implicated in BBS^17,26^.

### OligoPVP: Use of background knowledge to find causative combinations of variants

Our results demonstrate that PVP can identify combinations of variants implicated in a disease significantly outperforming current state-of-the-art gene prioritisation approaches. The variants found by PVP are commonly in genes that form a disease module, i.e., a set of interacting genes that are jointly associated with a disease or phenotype^27^. For example, out of the 165 di-allelic combinations used in our study, we can find evidence of interactions for 71 di-allelic combinations and 16 tri-allelic combinations using the interaction database STRING^25^. The STRING resource contains background knowledge about the interaction between genes based on protein-protein interactions, co-expression, pathway involvement, or co-mention in literature, and therefore provides a wide range of distinct interaction types which may underlie a phenotype. Using this background knowledge about which genes interact may be useful to further improve prioritization of variants in oligogenic diseases.

We have developed OligoPVP, an algorithm that uses background knowledge from interaction networks to prioritize variants in oligogenic diseases. OligoPVP identifies likely causative variants in interacting genes and ranks tuples of *n* variants in genes that are connected through an interaction network. OligoPVP will first rank all variants in a set of variants (such as those found in a VCF file) independently using PVP and assign each variant *v* a prediction score *σ*(*v*). When ranking combinations of *n* variants, OligoPVP will then evaluate all *n*-tuples of variants *v*_1_,…, *v_n_* and assign a score 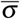 to the *n*-tuple (*v*_1_,…, *v_n_*), given an interaction network ϒ:

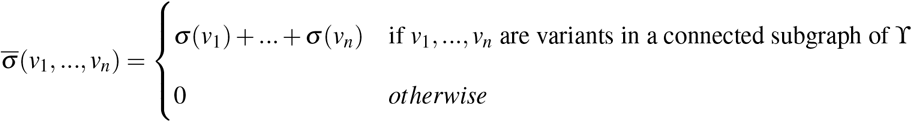

Algorithm 1 illustrates the procedure to find oligogenic disease modules in more detail. OligoPVP can identify combinations of variants both in exonic and non-exonic regions. For non-exonic variants, we assign the gene that is located closest to the variant as the variant’s gene.

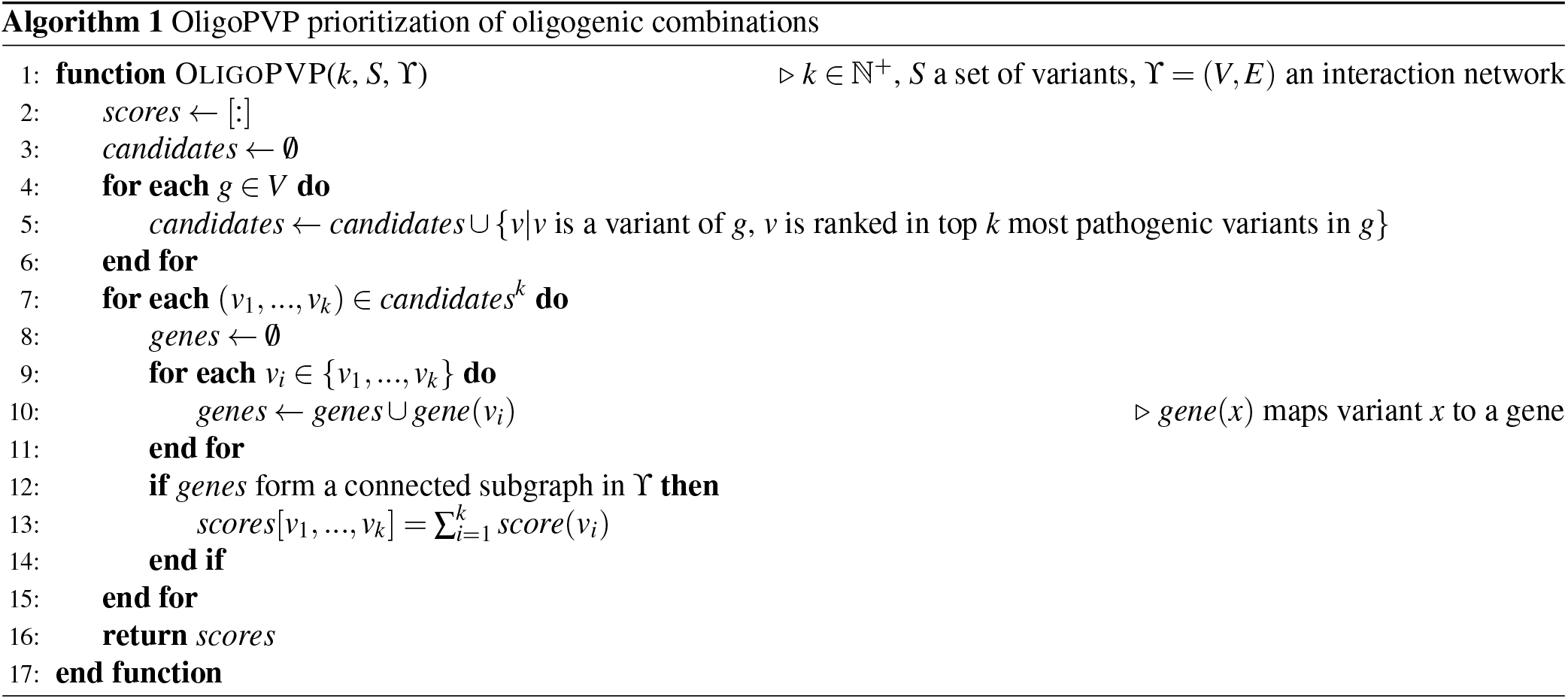

The OligoPVP algorithm strictly relies on an interaction network as background knowledge and will not prioritize any combinations of variants if they are not connected in an interaction network that is used as background knowledge. OligoPVP utilizes beam search^28^ to optimize memory usage. We can simply extend OligoPVP to also consider compound heterozygote combinations of variants by adding self-loops to each node in ϒ. The main advantage of OligoPVP is its ability to identify and rank connected sets of variants higher than individual variants. Table 4 lists several cases in which OligoPVP prioritizes pairs of variants significantly higher than PVP would prioritize them on their own. On the other hand, OligoPVP will not prioritize combinations of variants if they are in genes that are not connected in the background network ϒ. Supplementary Table 1 lists some of the cases which can be prioritized with PVP but not OligoPVP.

**Table 4.**
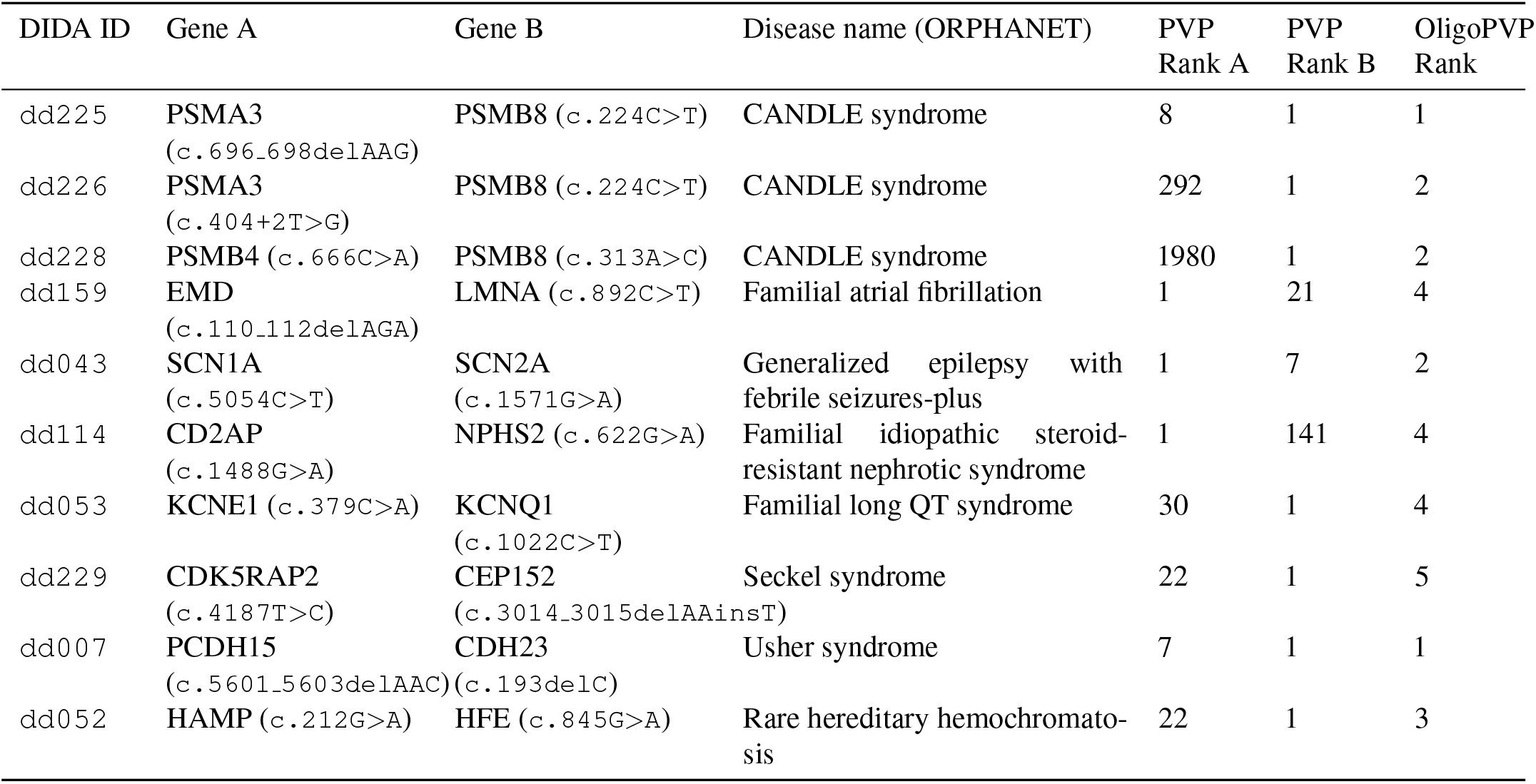
Cases of DIDA combinations improved by OligoPVP in comparison to PVP. OligoPVP incorporates protein-protein interactions in the prioritization of variant tuples. We compare the results of applying OligoPVP to the ranks obtained using PVP on individual variants.

| DIDA ID | Gene A | Gene B | Disease name (ORPHANET) | PVP Rank A | PVP Rank B | OligoPVP Rank |
| --- | --- | --- | --- | --- | --- | --- |
| dd225 | PSMA3<br>(c.696_698delAAG) | PSMB8 (c.224C>T) | CANDLE syndrome | 8 | 1 | 1 |
| dd226 | PSMA3<br>(c.404+2T>G) | PSMB8 (c.224C>T) | CANDLE syndrome | 292 | 1 | 2 |
| dd228 | PSMB4 (c.666C>A) | PSMB8 (c.313A>C) | CANDLE syndrome | 1980 | 1 | 2 |
| dd159 | EMD<br>(c.110_112delAGA) | LMNA (c.892C>T) | Familial atrial fibrillation | 1 | 21 | 4 |
| dd043 | SCN1A<br>(c.5054C>T) | SCN2A<br>(c.1571G>A) | Generalized epilepsy with febrile seizures-plus | 1 | 7 | 2 |
| dd114 | CD2AP<br>(c.1488G>A) | NPHS2 (c.622G>A) | Familial idiopathic steroid-resistant nephrotic syndrome | 1 | 141 | 4 |
| dd053 | KCNE1 (c.379C>A) | KCNQ1<br>(c.1022C>T) | Familial long QT syndrome | 30 | 1 | 4 |
| dd229 | CDK5RAP2<br>(c.4187T>C) | CEP152<br>(c.3014_3015delAAinsT) | Seckel syndrome | 22 | 1 | 5 |
| dd007 | PCDH15<br>(c.5601_5603delAAC) | CDH23<br>(c.193delC) | Usher syndrome | 7 | 1 | 1 |
| dd052 | HAMP (c.212G>A) | HFE (c.845G>A) | Rare hereditary hemochromatosis | 22 | 1 | 3 |

## Discussion

With the increasing appreciation of the relationship between complex and Mendelian diseases^29^, the ability to discover multiple contributing variants in the same genome provides a powerful tool to help understand the genetic architecture of diseases and the underlying physiological pathways. With the advent of whole exome and whole genome sequencing, advances have been made using existing approaches to prioritize causative variants. However, use of standard criteria for the identification of rare disease variants, e.g., a low minor allele frequence (MAF) of, for example, less than 1%, are designed to detect *de novo*, homozygous, or compound heterozygous variants, and may not give sufficient priority to variants of low apparent pathogenicity, haploinsufficiency, or low to medium MAF, although these variants may still be important in the pathogenesis of a disease. Because the approach we take with OligoPVP and PVP makes no assumptions about allele frequency or mode of inheritance, and balances estimates of pathogenicity with phenotypic relatedness, a wider net is cast and candidate genes affecting the penetrance, expressivity or spectrum of the phenotype are more readily identified.

Genes whose variants contribute to a disease phenotype are considered likely to sit within the same pathway or network^30–33^. In addition to well established studies of genes involved in, for example the ciliopathies^26,34^, newer studies are now identifying network relations between genes implicated in the oligogenic origins of diseases^14,35^. Consequently, we can exploit background knowledge on the interactions of gene products in OligoPVP and improve the ranking of candidate pairs of variants over that assigned through pathogenicity and phenotypic relatedness scores alone.

Currently, identification of multiple variants contributing to the characteristics of a disease in a cohort or individual patient rely either on a candidate gene approach or the assumption that contributing alleles are likely to be rare in the population. The contribution of rare alleles of low effect, i.e., which by themselves generate sub-clinical phenotypes, for example hypomorphs, may be missed in this way, and rare to medium frequency alleles which modify the penetrance or expressivity of a second remain difficult to identify (the former because of low potential pathogenicity and the latter because of high frequency and lack of association with a phenotype when occurring alone). An alternative strategy for identification of candidate genes for highly heterogeneous human diseases is to use mouse genetics to identify phenotypic modifier genes. For example, neural tube defects are believed to involve more than 300 genes in the mouse, mutations in many of which need to be digenic or trigenic for expression of the phenotype^36^. The scale of genetic interactions becoming apparent from mouse studies strongly supports the suggestion that in the human, we are only seeing the tip of a very important iceberg^37^.

The OligoPVP algorithm aims to present a generic framework for using background knowledge about any form of interaction between genes and gene products to guide the identification of combinations of variants. In its generic form, the worst case complexity of the algorithm is 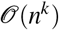 where *n* is the number of variants and *k* the size of the module (the size of the module is a parameter of OligoPVP). It is clear that our algorithm, in its basic form, will not yet scale to large disease modules (i.e., large *k*); however, in the future, several methods can be used to further improve the average case complexity to find larger disease modules.

Furthermore, background knowledge about interactions between genes and gene products is far from complete. In particular, information about coarse scale physiological interactions, i.e., those that occur based on whole organism physiology, are significantly underrepresented in interaction databases^19^. Additionally, interaction networks may have biases such as overrepresentation of commonly studied genes^38,39^, and these biases will likely effect the performance of our algorithm. As more genomic data related to complex diseases becomes available, more work will be required to identify and remove these biases in the identification of phenotype modules from personal genomic data.

OligoPVP is, to the best of our knowledge, the first phenotype-based method to identify disease modules in personal genomic data. With the large (i.e., exponential) number of combinations of variants that have to be evaluated in finding disease modules, it is clear that any computational method has to make use of background knowledge to restrict the search space of potentially causative combinations of variants. OligoPVP is such a method which uses knowledge about interactions and phenotype associations to limit the search space. In the future, more background knowledge can be incorporated to improve OligoPVP’s coverage as well as accuracy. OligoPVP is freely available at https://github.com/bio–ontology–research–group/phenomenet–vp.

## Acknowledgements (not compulsory)

This work was supported by funding from King Abdullah University of Science and Technology (KAUST) Office of Sponsored Research (OSR) under Award No. URF/1/3454-01-01 and FCC/1/1976-08-01. GVG acknowledges support from H2020-EINFRA (731075) and the National Science Foundation (IOS:1340112) as well as support from the NIHR Birmingham ECMC, NIHR Birmingham SRMRC and the NIHR Birmingham Biomedical Research Centre and the MRC HDR UK. The views expressed in this publication are those of the authors and not necessarily those of the NHS, the National Institute for Health Research, the Medical Research Council or the Department of Health.

